# Powerful statistical method to detect disease associated genes using publicly available GWAS summary data

**DOI:** 10.1101/478321

**Authors:** Jianjun Zhang, Zihan Zhao, Xuan Guo, Bin Guo, Baolin Wu

## Abstract

Genome-wide association studies (GWAS) have thus far achieved substantial success. In the last decade a large number of common variants underlying complex diseases have been identified through GWAS. In most existing GWAS, the identified common variants are obtained by single marker based tests, that is, testing one single nucleotide polymorphisms (SNP) at a time. Generally the basic functional unit of inheritance is a gene, rather than a SNP. Thus, results from gene level association test can be more readily integrated with downstream functional and pathogenic investigation. In this paper, we propose a general gene-based p-value adaptive combination approach (GPA) which can integrate association evidence of multiple genetic variants using only GWAS summary statistics (either p-value or other test statistics). The proposed method could be used to test both continuous and binary traits through not only a single but also multiple studies, which helps overcome the limitation of existing methods that only can be applied to specific type of data. We conducted thorough simulation studies to verify that the proposed method controls type I errors well, and performs favorably compared to single-marker analysis and other existing methods. We demonstrated the utility of our proposed method through analysis of GWAS meta-analysis results for fasting glucose and lipids from the international MAGIC consortium and Global Lipids Consortium, respectively. The proposed method identified some novel traits associated genes which can improve our understanding of the mechanisms involved in *β*-cell function, glucose homeostasis and lipids traits.

## Introduction

Genome-wide association studies (GWAS) have been remarkably successful in identifying large number of genetic variants associated with complex traits and diseases. However, these identified genetic variants can only explain a small to modest fraction of the overall heritability for most complex traits and diseases (Manolio et al., 2009). This partially because most GWAS are primarily based on the paradigm of single-variant single-trait association tests without considering the Single Nucleotide Polymorphism (SNP) dependence in genes. Combining multiple SNPs to conduct SNP set-based or gene-based association tests can increase power compared to the single SNP based tests (Wu et al., 2010). There have been some set-based or gene-based association test methods which have demonstrated their utility to boost detection power and identify more genetic variants, however, these methods generally require raw genotype and phenotype data. Due to privacy concerns and various logistic considerations, raw individual-level phenotype and genotype data are usually very difficult to obtain. Although current techniques can sequence the whole genome of large groups of individuals for all genetic variants, they are not very cost effective. It has been a common practice that many GWAS make their summary statistics results available to the public. These summary data generally include the minor allele frequency (MAF), the test statistics, and significance p-values for each SNP. The publicly available GWAS data and the motivation to identify more traits associated genes to explain the missing heritability inspired us to develop novel method by integrating the results from standard single variant analysis using the GWAS summary data (Pasaniuc and Price, 2017).

Recently several statistical association test methods have been developed using only GWAS summary data. Bakshi et al. (2016) proposed a method that can test SNP-set association with a single trait using GWAS summary data. Pasaniuc and Price (2017) reviewed recent progress on statistical methods that leverage GWAS summary data to gain insights into the genetic basis of complex traits and diseases. Stephens (2013) and Zhu et al. (2015) observed that the correlation matrix of test statistics from GWAS summary data is the same as the correlation of traits for a single variant. Based on this finding they proposed statistical methods to test association between a single variant and multiple correlated traits using individual GWAS summary statistics and GWAS meta-analysis summary results. Guo and Wu (2018) proposed three SNP-set association tests: sum test (ST), squared sum test (S2T) and adaptive test (AT). The sum test is a type of burden test statistic that can perform well when all variants have the same direction of effects, but it may suffer low statistical power when the effects are in different directions. The squared sum test is a type of quadratic sum test that could be less powerful when we consider a large number of SNPs simultaneously, although it performs better for SNP-sets with both protective and risk variants. The third adaptive test combined the strength of the sum test and squared test by considering their optimal combination and thus may lose some information from other combinations. Note that these existing methods are only built on the summary Z-statistics, which will have some restrictions when some GWAS only report p-values or other types of summary test statistics. In addition, these test statistics results may be computed using the same or different statistical methods. The traits of interest could be independent or correlated and continuous or binary, and they may be analyzed through a single study or multiple studies. It is desirable to develop powerful and generalized association test methods which could be used for different types of GWAS summary data.

Motivated by the aforementioned considerations, in this paper, we propose a general gene-based p-value adaptive combination approach that can integrate association evidence from GWAS summary statistics, either p-values or other statistical values, for continuous or binary traits, which might come from the same or different studies of traits. In order to increase power for the gene-based association test, our method searches for a strongly associated subset of SNPs by ranking p-values to identify strong disease associated genes. We show that the correlation of the Z-statistics across variants can be computed based on the variant linkage disequilibrium (LD) matrix. Furthermore, we show that at the permutation steps to evaluate p-value of our proposed adaptive method, we can leverage LD information to estimate the correlation of Z-statistics. We develop algorithms for the proposed method which can compute p-values very efficiently and accurately. The proposed method is very useful to mine the vast amount of public GWAS summary data to identify additional interesting genetic variants. Results obtained from analyzing GWAS meta-analysis summary results of fasting glucose and lipids suggests that the proposed method has improved statistical power over single-marker analysis and other existing methods.

## Methods

Consider a sample of *n* unrelated individuals for a continuous trait and each individual has been genotyped at *M* variants in a genomic region (a gene or a pathway). Let *Y* = (*y*_1_, *…, y*_*n*_)^*T*^ denote the phenotype of *n* individuals, *G*_*m*_ = (*g*_1*m*_, *…, g*_*nm*_)^*T*^ denote the *m*^*th*^ genotypic score of *n* individuals, where *g*_*im*_ ∈ {0, 1, 2} is the number of minor alleles that the *i*^*th*^ individual carries at the *m*^*th*^ genetic variant.

We first consider a simplified case without covariates and describe the relationship between the phenotype *Y* and the genotypic score of the *m*^*th*^ variant *G*_*m*_ using linear regression model:

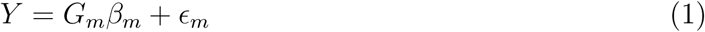

where *∈*_*m*_ is assumed to follow a normal distribution with mean zero and variance 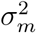. Naturally, we have :

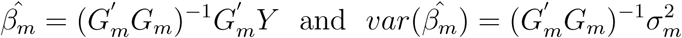

If we consider Z-statistics for GWAS, the corresponding Z-statistics are computed as the estimated regression coefficients of the genetic terms divided by their estimated standard errors, written as:

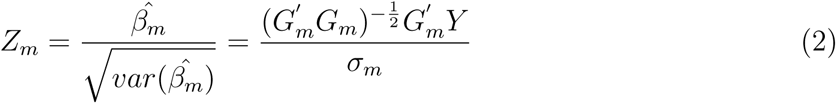

Even though the variance 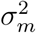 should vary across different SNPs in general, 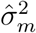s are all equal and they are unbiased estimates of variance 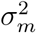 for the same continuous phenotype based on mode (1) under null hypothesis *β*_1_ = … = *β*_*M*_ = 0. We can easily check that *var*(*Z*_*m*_) = 1 for *m* = 1, *…, M* and

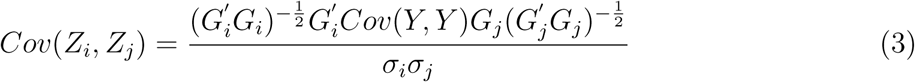

based on equation (2). When the unknown variance 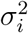 or 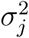 is replaced by their estimate 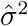 under null hypothesis, the formula (3) indicate that:

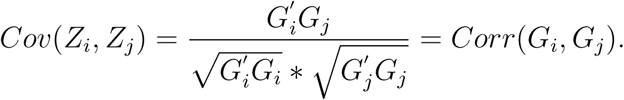

Under general case with covariates, we can also obtain the same result when the genotypes of genetic variants are independent with covariates, referred to Guo and Wu (2018) using projection matrix. It means that the covariance or correlation between summary Z-statistics equals to the LD correlation matrix of variants’ genotypes. The LD matrix (Hereafter denoted as *R*) can often be computed from publicly available samples for relatively homogeneous population.

Based on summary statistics ***p*** = (*p*_1_, *…, p*_*M*_)*′* of a set of *M* variants in a gene region, we propose a new method, motivated by Liang et al. (2016), to test the null hypothesis *H*_0_: there is no association between the continuous trait and the *M* variants. Without loss of generality, we assume that the p-values are based on the Z-statistics, denoted as ***Z***. It is reasonable to assume that ***Z*** follows a multivariate normal distribution with mean 0 and correlation matrix R under the null hypothesis. Let *p*_1_, *p*_2_, *…, p*_*M*_ denote the corresponding values for the *M* variants. Based on these p-values, we propose a general gene based p-value adaptive combination approach (GPA) to test the association between the continuous trait and these *M* genetic variants where the statistic of GPA is written as:

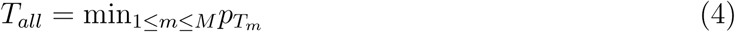

where *p*_*T*_*m* denotes the p-value of summary statistic 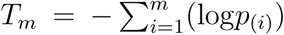, in which *p*_(*i*)_ denotes the *i*^*th*^ smallest p-value of *p*_1_, *p*_2_, *…, p*_*M*_. Without needing to generate genotype and phenotype data, we use the following permutation procedure to evaluate the p-values of our statistic *T*_*all*_ by simulating the Z-statistic from multivariate normal distribution *N* (0, *R*):

1. In each permutation, based on ***Z*** *∼ N* (0, *R*), we recalculate *p*_(1)_,*…, p*_(*M*)_ and *T*_1_,*…, T*_*M*_. Suppose that we perform B times of permutations. Let 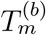, *b* = 1,…, *B* denote the value of *T*_*m*_ based on the *b*^*th*^ permuted data, where *b* = 0 represents the value computed by the original summary data.
2. For the *m*^*th*^ genetic variant, we transfer 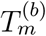 to 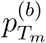 by:

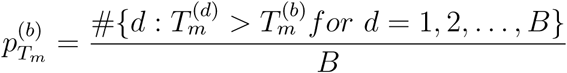
3. Let 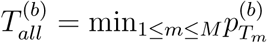, then the p-value of *T*_*all*_ is given by:

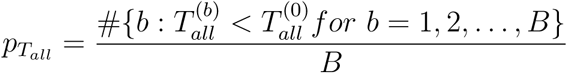

Because the null distribution of ***Z*** = (*Z*_1_, *…, Z*_*M*_) follows multivariate normal distribution with means zero and covariance matrix *R*. The distribution *p*_1_, *p*_2_, *…, p*_*M*_ and *T*_*all*_ only depend on the correlation between genetic variants and don’t depend on trait values, thus the permutation procedure described above to generate an empirical null distribution of *T*_*all*_ needs to be done only once.

## Results

### Comparison of Methods

Currently, there are three types of methods utilizing summary statistics including burden, quadratic and adaptive forms (Pasaniuc and Price, 2017). The three methods proposed by Guo and Wu (2018): sum test (ST), squared sum test (S2T) and adaptive test (AT) are good representative of the three types of methods. Therefore, we compare the performance of the proposed method (GPA) with the three methods. Here we briefly introduce each of these three methods using the notations in the Methods section.

1. Sum test (ST), 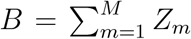, a type of burden test statistic (Madsen and Browning, 2009).
2. Squared sum test (S2T), 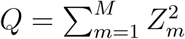, a type of SKAT statistic (Wu et al., 2010).
3. Adaptive test (AT), *T* = min_*ρ∈*[0,1]_ *P*(*Q*_*ρ*_), where *Q*_*ρ*_ = (1 − *ρ*)*Q* + *ρB*^2^ and *P* (*Q*_*ρ*_) denote the corresponding p-value.

Because 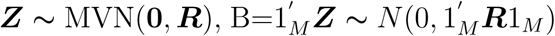 where 1_*M*_ denote column vectors of length *M* that are all 1’s. That is, 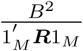 follows the *χ*_1_ distribution. Q=***Z****′* ***Z*** is asymptotically distributed as the weighted sum of independent *χ*_1_ random variables with weights being the eigen-values of ***R***. The p-value of T can be efficiently and accurately computed by an one-dimensional numerical integration where we will search over *ρ ∈* (0, 0.01, 0.04, 0.09, 0.16, 0.25, 0.5, 1) following Wu et al. (2016). The p-value of these three statistics can be obtained using the “sats” function in the “mkatr” package in R.

In addition, we compare the GPA method with the minimum p-value approach, which takes the minimum p-value across all SNPs in the test gene.

## Simulations

### Type I error

We conducted simulation studies to evaluate the performance of the proposed method with the three comparable methods. For the type I error evaluation, instead of generating genotype and phenotype data, we simulate the test statistic ***Z*** from a multivariate normal distribution *N* (**0**, *R*) where *R* is the LD correlation matrix of gene NPHS2 including 20 SNPs reported in Dupuis et al. (2010) study. We consider four different significance levels *α* = 5 *×* 10^−2^, 10^−2^, 10^−3^, 10^−4^. The estimated type I error rates are summarized in Table 1. In this simulation, p-values are estimated by performing 10^5^ times permutations and over 10^5^ replicates. From this table, we can see that all estimated type I error rates are not significantly different from the nominal levels and thus these tests are all valid tests.

**Table 1:** Estimated type I error rates for ST, S2T, AT and our developed method GPA.

| α-level | ST | S2T | AT | GPA |
| --- | --- | --- | --- | --- |
| $5 \times 10^{-2}$ | 0.051 | 0.050 | 0.053 | 0.052 |
| $1 \times 10^{-2}$ | 0.006 | 0.010 | 0.011 | 0.009 |
| $1 \times 10^{-3}$ | 0.001 | 0.001 | 0.0008 | 0.0012 |
| $1 \times 10^{-4}$ | 0.0001 | 0.0001 | 0.00009 | 0.0001 |

### Power comparison

To evaluate the power of the proposed method, we conduct simulations using gene NPHS2 with 20 SNPs. Consider for the comparability of these methods and avoiding time-consuming, we simulate 10^3^ summary statistics from *N* (***A***, *R*) where ***A*** is a vector of length 20. We consider nine scenarios: assuming 6 SNPs with association signals in the first seven scenarios and 4 SNPs with association signals in the remaining two scenarios and setting the rest elements of ***A*** as zero. In order to make fair and compatible comparison of these methods, we assign appropriate effect size for causal SNPs. *R* is the LD correlation matrix of the gene NPHS2. In each permutation, p-values are estimated by performing 10^4^ times permutations. Table 2 shows the estimated power at 10^−3^ significance level under different settings of ***A***. Overall the proposed GPA test performs robustly across all scenarios and has the overall best performance comparing with the other three tests. Among the four tests, when there are different directions of effects across all 4 signal SNPs, the burden test ST has the worst performance and the adaptive method AT also suffer loss of power. The squared sum test (S2T) loses power because of the effect of the other noises. Our proposed test GPA is robust to different directions of effects and effect of the other noises.

**Table 2:**
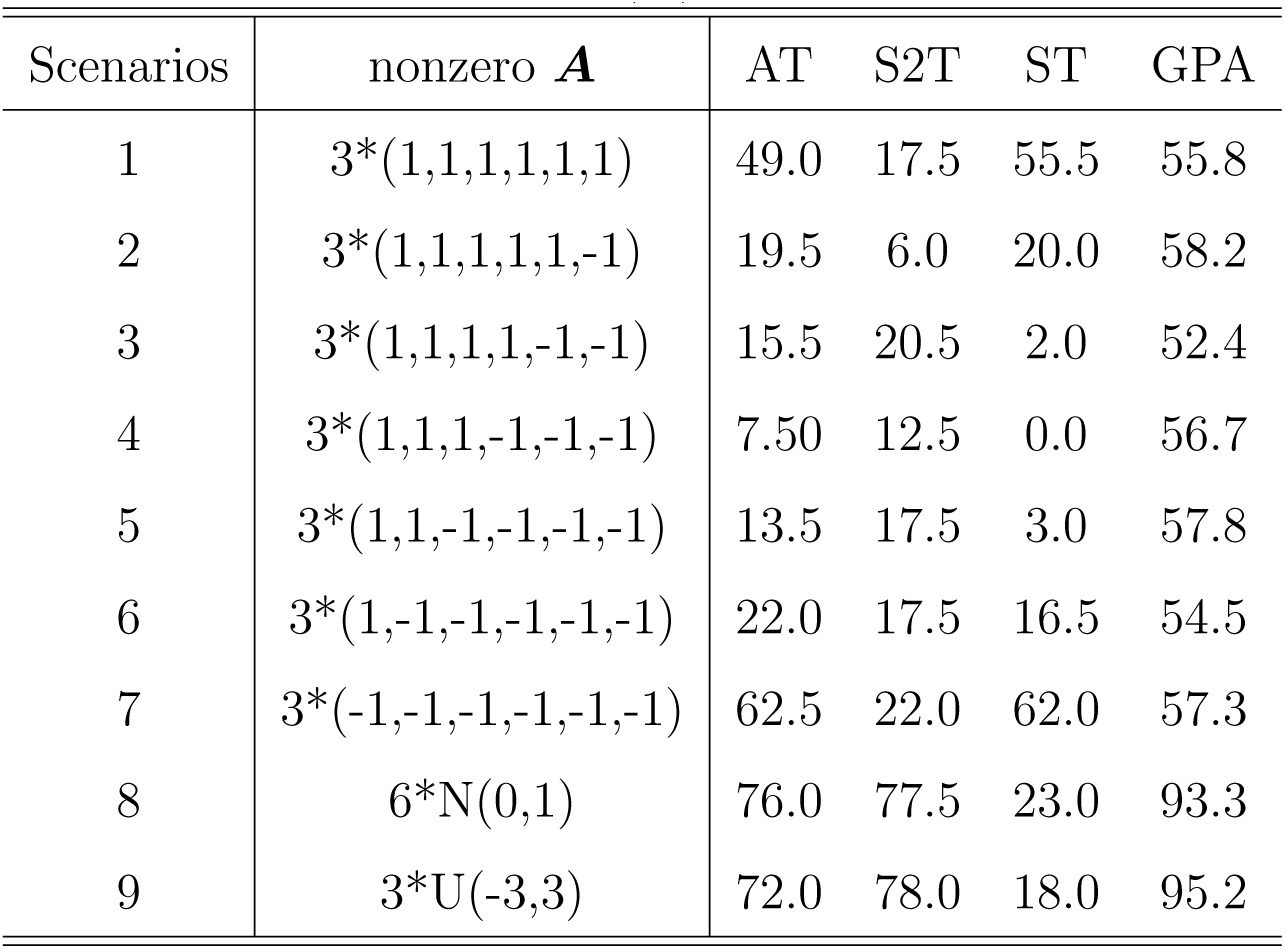
Estimated test power (%) under 10^−3^ significance level.

## Real data analysis

### Application to fasting glucose GWAS meta-analysis summary results

The summary data of fasting glucose from GWAS meta-analysis was conducted by the international MAGIC consortium (Dupuis et al. 2010), which are based on 21 GWAS with 46,186 non-diabetic participants of European descent who are informative for fasting glucose. It consists of the MAF, effect size estimate and its associated standard error, Z-statistic, and p-value for 18,725 genes containing 2,470,476 SNPs. The summary data can be found from ftp://ftp.sanger.ac.uk/pub/magic/MAGIC_FastingGlucose.txt. We download the list of genes and their coordinates (transcription start and end positions based on the hg19/GRChB37 reference genome) from the UCSC genome browser (Kent et al., 2002). We take all SNPs that are located in or near a gene as a set to be analyzed for joint association and group all SNPs from 20 kb upstream of a gene to 20 kb downstream of a gene following Wu et al. (2010).

We use another summary data of Dupuis et al. (2010) as partial validation for our analysis. The summary data is performed by MAGIC consortium for a followup replication study using a Metabochip consisting of a small panel of promising SNPs and a much larger sample size from 66 fasting glucose GWAS with around 133,010 non-diabetic participants (Scott et al. 2012). The summary data contains the similar results for 64,493 pre-selected SNPs which is available at ftp://ftp.sanger.ac.uk/pub/magic/MAGIC_Metabochip_Public_data_release_25Jan.zip.

In order to better illustrate the GPA method, we remove 290 genome-wide significant SNPs with p-value less than 5 *×* 10^−8^, filter out those SNPs with MAF*<* 0.05 and remove SNPs that have pairwise LD *r*^2^ *>* 0.8 with other SNPs. For the 18,725 SNP sets, we set our genome-wide SNP set significance level as 0.05*/*18725 = 2.67 *×* 10^−6^, which is the Bonferroni corrected significance level for the total number of tested SNP sets. P-values of GPA are estimated by performing 10^7^ times permutations.

Figure 1 shows the Venn diagram comparing the number of significant genes identified by the proposed test with Guo and Wu’s method. The three methods of Guo and Wu (2018) identified 25 significant genes in total, specifically, the ST identified 12 significant genes, S2T identified 20 genes, and AT identified 22 significant genes, and our new method GPA identified 26 genes. Our method captured the majority of significant SNP-sets identified by S2T, ST and AT, and it further identified ten additional significant genes which were not identified by these three comparable methods. Table 3 shows the four test p-values for the ten genes and the minimum p-value across all SNPs in the gene from Dupuis et al. (2010) and Scott et al. (2012). All of the ten genes harbored genome-wide significant SNPs at Dupuis et al. (2010) study.

**Table 3:** Significant genes were identified only by GPA, missed by S2T, ST and AT: we listed these four test p-values and the minimum p-value across all SNPs in the the gene from Dupuis et al. (2010) study (denoted as minP-2010) and Scott et al. (2012) study (denoted as minP-2012).

| Gene | S2T | ST | AT | GPA | minP-2010 | minP-2012 |
| --- | --- | --- | --- | --- | --- | --- |
| MYL7 | 2.42E-05 | 8.70E-05 | 1.70E-05 | <5.00E-08 | 2.32E-12 | 4.22E-03 |
| G6PC2 | 6.66E-05 | 5.03E-04 | 7.55E-05 | <5.00E-08 | 4.61E-75 | 2.22E-02 |
| FEN1 | 1.27E-05 | 1.68E-01 | 1.87E-05 | 5.00E-07 | 3.82E-09 | 1.04E-01 |
| SPC25 | 3.90E-06 | 8.30E-03 | 8.55E-06 | <5.00E-08 | 4.61E-75 | 4.61E-03 |
| SLC35C1 | 2.91E-04 | 1.09E-03 | 3.78E-04 | <5.00E-08 | 1.18E-09 | 1.86E-02 |
| MIR611 | 1.27E-05 | 1.68E-01 | 1.87E-05 | <5.00E-08 | 3.82E-09 | 1.05E-1 |
| C11orf10 | 1.05E-04 | 3.19E-01 | 2.61E-04 | <5.00E-08 | 3.82E-09 | 1.05E-1 |
| MRPL33 | 3.67E-04 | 8.07E-02 | 6.67E-04 | <5.00E-08 | 1.25E-08 | 7.32E-03 |
| ABCB11 | 2.17E-05 | 4.92E-05 | 1.26E-05 | <5.00E-08 | 4.61E-75 | 5.4E-02 |
| TCF7L2 | 1.80E-05 | 8.00E-02 | 3.49E-05 | <5.00E-08 | 1.24E-08 | 3.48E-18 |

**Figure 1.**
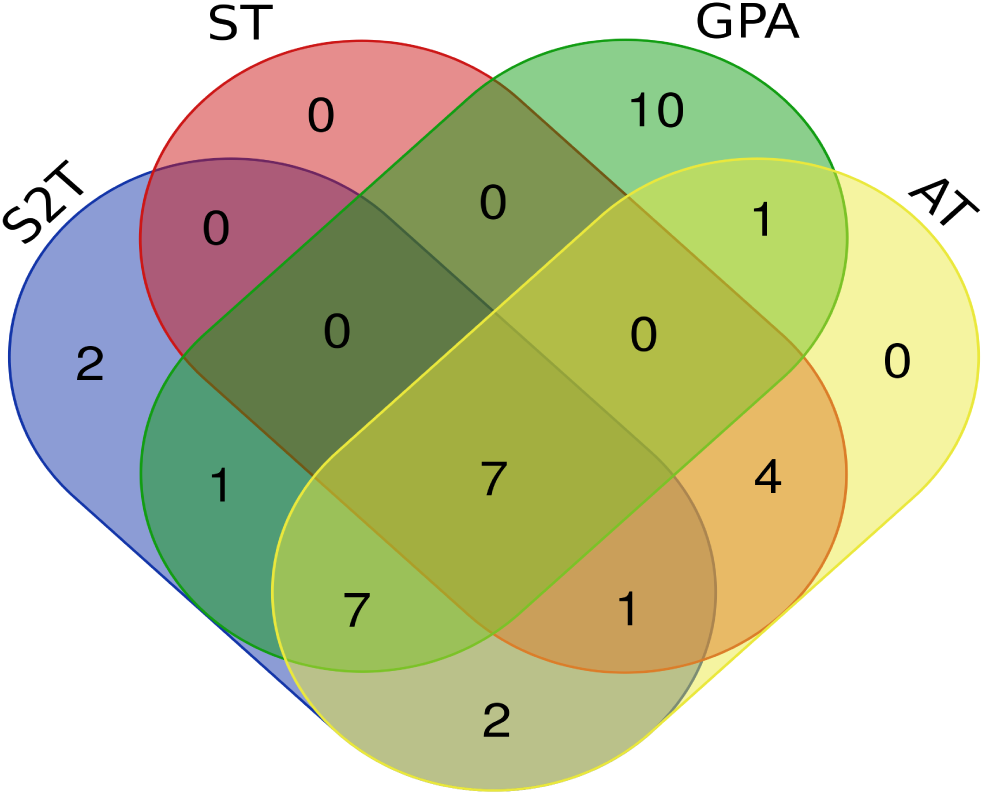
Venn diagram of number of significant genes identified by our method GPA with the methods of Guo and Wu (2018) for fasting glucose.

Figure 2 shows the Venn diagram comparing the number of significant genes identified by the proposed test with the minimum p-value across all SNPs in the gene from Dupuis et al. (2010) and Scott et al. (2012). In total we identified 2 novel genes: SLC4A1AP and SUPT7L. They did not harbor any significant SNPs in Dupuis et al. (2010) and Scott et al. (2012). Table 4 summarizes the information of the two genes. For each novel gene, we listed the p-values of the GPA test together with Guo and Wu’s tests and minimum p-value of all SNPs in the considered gene for both studies.

**Table 4:** GPA test p-values for 2 novel genes: we also listed these four test p-values and the minimum p-value across all SNPs in the gene from Dupuis et al. (2010) study (denoted as minP-2010) and Scott et al. (2012) study (denoted as minP-2012).

| Gene | S2T | ST | AT | GPA | minP-2010 | minP-2012 |
| --- | --- | --- | --- | --- | --- | --- |
| SLC4A1AP | 1.22E-06 | 6.06E-01 | 1.51E-06 | <5.00E-08 | 2.96E-6 | 5.10E-02 |
| SUPT7L | 1.76E-06 | 4.15E-04 | 2.24E-06 | <5.00E-08 | 2.96E-06 | 6.80E-02 |

**Figure 2.**
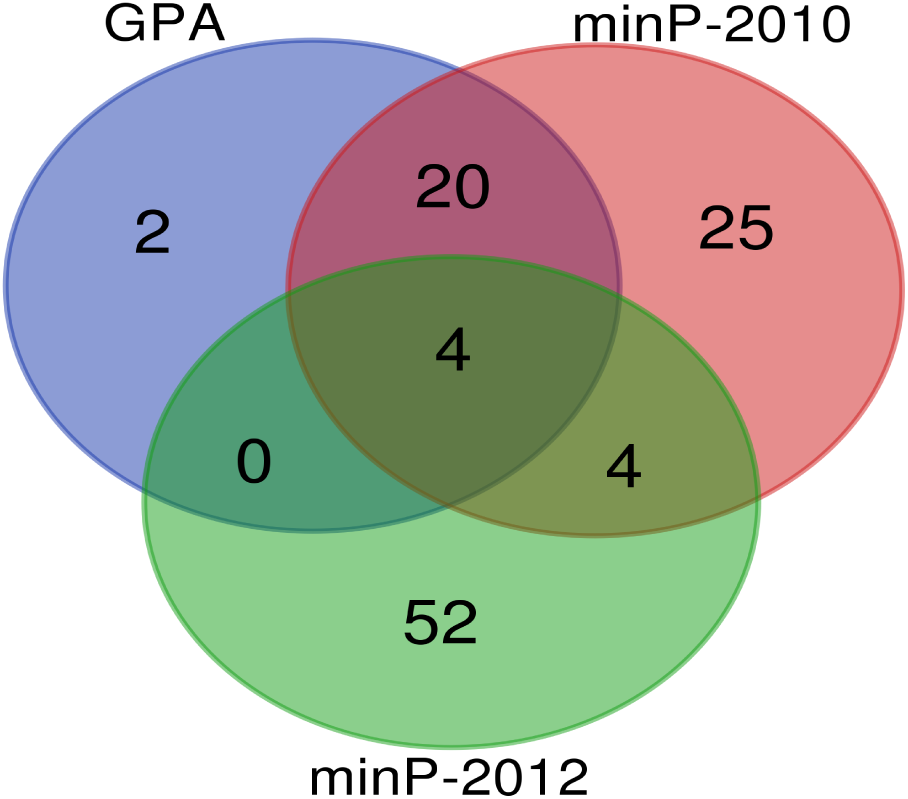
Venn diagram of number of significant genes identified by our method GPA with minP-2010 and minP-2012 for fasting glucose.

### Application to GWAS summary data for lipids traits

We also conduct a comprehensive analysis of the GWAS summary data for high-density lipoprotein cholesterol (HDL), low-density lipoprotein cholesterol (LDL), and Triglycerides (TG) conducted by the Global Lipids Consortium with around 100, 000 European individuals (Teslovich et al., 2010). The GWAS summary data are downloaded from http://csg.sph.umich.edu/ abecasis/public/lipids2010. We follow the previous procedure to perform the summary data. Willer et al.(2013) conducted a followup study of those promising SNPs identified by Teslovich et al. (2010) based on around 190,000 European participants. We use their summary results as partial validation in our analysis. Here we use the minimum p-values across these three traits for these four test methods to test significant genes.

Figure 3 shows the Venn diagram comparing the number of significant genes identified by the proposed tests with Guo and Wu’s method. At the Bonferroni corrected SNP-set significance level 2.67 *×* 10^−6^, our proposed GPA test identified a total of 302 significant genes and the three methods of Guo and Wu (2018) identified 254 significant genes in total. Specifically, the ST identified 108 significant genes, S2T identified 206 genes, and AT identified 234 significant genes. Obviously, our method GPA captured the majority of significant genes identified by S2T, ST and AT, and it further identified 92 additional significant genes which were not identified by these three comparable methods. Among the 92 significant genes, 90 genes harbored genome-wide significant SNPs at Teslovich et al. (2010) study. Table 5 shows 10 representative genes of these 92 genes by these four tests together with the minimum p-value across all SNPs in the gene from Teslovich et al. (2010) and Willer et al. (2013).

**Table 5:** Ten significant genes were identified only by GPA, missed by S2T, ST and AT: we listed the p-values of the four tests and the minimum p-value across all SNPs in the gene from Teslovich et al. (2010) study (denoted as minP-2010) and Willer et al. (2013) study (denoted as minP-2013).

| Gene | S2T | ST | AT | GPA | minP-2010 | minP-2013 |
| --- | --- | --- | --- | --- | --- | --- |
| HSD3B7 | 2.09E-05 | 8.47E-06 | 1.19E-05 | <5.00E-08 | 1.51E-07 | 1.03E-07 |
| BAIAP2L2 | 9.82E-06 | 4.89E-01 | 2.76E-05 | <5.00E-08 | 2.45E-07 | 2.50E-07 |
| ACSS2 | 5.62E-05 | 4.39E-02 | 1.10E-04 | <5.00E-08 | 7.83E-08 | 8.50E-07 |
| ASGR1 | 2.02E-05 | 1.40E-02 | 4.03E-05 | <5.00E-08 | 5.40E-07 | 2.81E-10 |
| FRMD5 | 5.30E-05 | 1.21E-01 | 6.42E-05 | <5.00E-08 | 1.63E-11 | 4.84E-17 |
| ZSWIM1 | 6.49E-06 | 4.94E-04 | 6.02E-06 | <5.00E-08 | 6.57E-08 | 5.21E-08 |
| MYBPC3 | 4.27E-06 | 2.34E-04 | 4.75E-06 | <5.00E-08 | 6.18E-16 | 3.30E-38 |
| SF4 | 6.42E-05 | 3.97E-03 | 1.40E-04 | <5.00E-08 | 2.90E-38 | 4.12E-77 |
| STX1B | 1.95E-05 | 3.52E-05 | 1.11E-05 | <5.00E-08 | 1.51E-07 | 1.03E-07 |
| GNAI3 | 4.68E-06 | 6.86E-03 | 8.46E-06 | <5.00E-08 | 6.35E-08 | 5.60E-14 |

**Figure 3.**
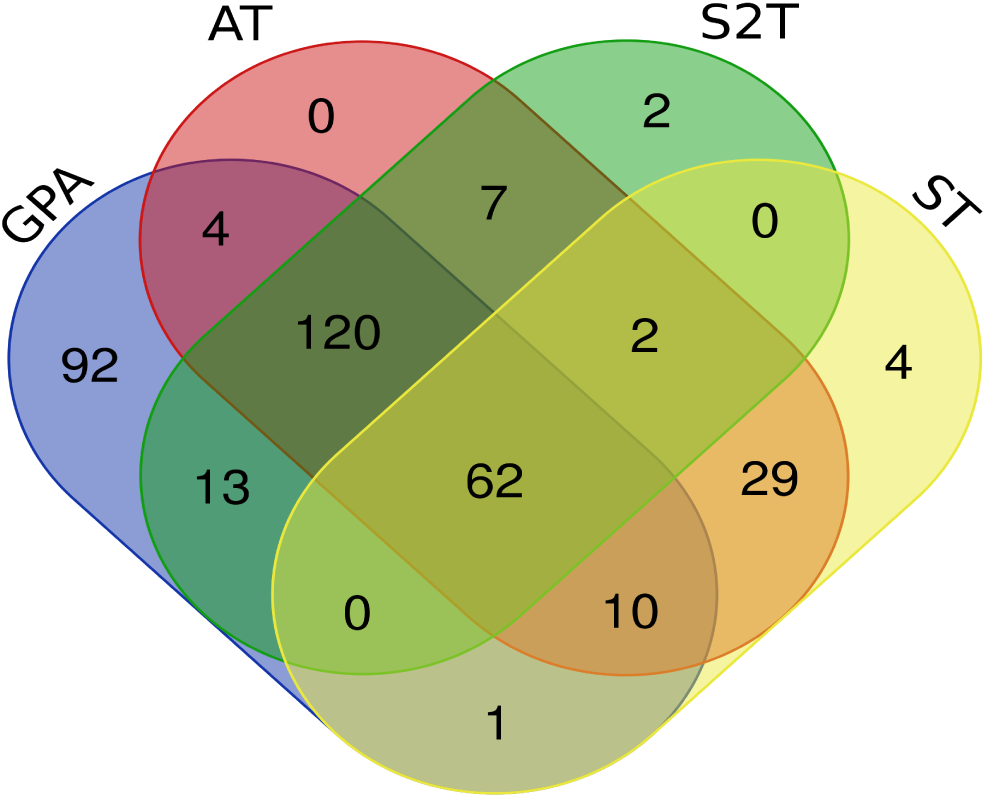
Venn diagram of the number of significant genes identified by our method GPA and the methods of Guo and Wu (2018) for lipids traits.

Figure 4 shows the Venn diagram comparing the number of significant genes identified by the proposed test with the minimum p-value across all SNPs in the gene from Teslovich et al. (2010) and Willer et al. (2013). Our proposed method identified 5 novel significant genes. Table 6 summarizes the five novel genes identified by GPA test. It shows the GPA test p-values together with minimum p-value of all SNPs in the gene across all traits for the two studies: Willer et al. (2013) and Teslovich et al. (2010). Among the 302 significant genes identified by our method, 299 genes harbored genome-wide significant SNPs from the meta-analysis results of Willer et al. (2013); and 5 of these 303 significant genes were novel genes in the sense that the meta-analysis of Teslovich et al. (2010) did not identify any significant SNPs in these 5 genes. Interestingly, gene PTPN13 was only identified by our proposed method GPA.

**Table 6:** GPA test p-values for 5 novel genes: we listed the four tests p-values and the minimum p-value across all SNPs in the gene from Teslovich et al. (2010) study (denoted as minP-2010) and Willer et al. (2013) study (denoted as minP-2013).

| Gene | S2T | ST | AT | GPA | minP-2010 | minP-2013 |
| --- | --- | --- | --- | --- | --- | --- |
| PTPN13 | 1.31E-05 | 1.03E-02 | 2.19E-05 | <5.00E-08 | 3.41E-05 | 8.17E-06 |
| POM121B | 4.45E-06 | 5.69E-09 | 2.37E-08 | <5.00E-08 | 9.58E-05 | 5.95E-07 |
| CPNE2 | 9.13E-07 | 5.18E-06 | 1.59E-06 | <5.00E-08 | 8.27E-06 | 7.41E-10 |
| MAFB | 3.88E-06 | 3.37E-02 | 9.26E-06 | 1.10E-06 | 1.78E-05 | 4.65E-07 |
| NSUN5 | 4.45E-06 | 5.69E-09 | 2.38E-08 | <5.00E-08 | 9.58E-05 | 5.95E-07 |

**Figure 4.**
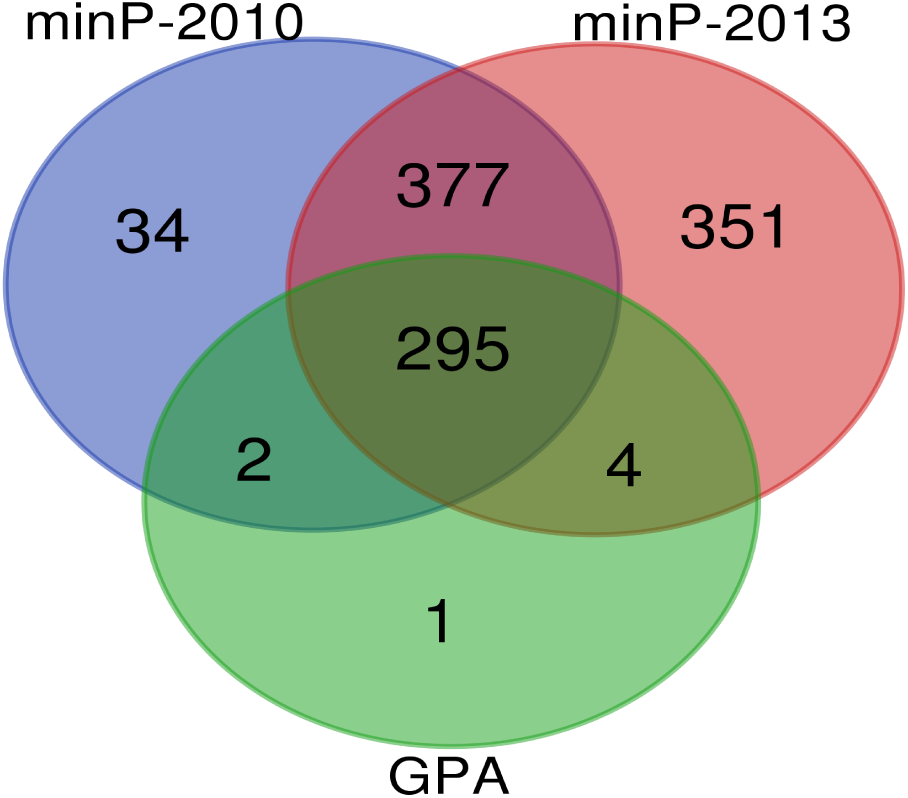
Venn diagram of the number of significant genes identified by our method GPA and minP-2010 and minP-2013 for lipids traits.

### Summary

We summarized our findings from fast glucose and lipids traits based on all the real data (Dupuis et al. 2010, Scott et al. 2012, Teslovich et al. 2010 and Willer et al. 2013). Two genes (SLC4A1AP, SUPT7L) and one gene (PTPN13) are only identified by our proposed method for these two studies, respectively. These genes have been reported to be significantly associated with fast glucose and lipids traits or some related traits which are strongly related to the fast glucose and lipids traits. For example, coronary artery disease (CAD) is a common disease showing strong relationship with lipids traits (Nair et al., 2009; Wilson, 1990). High blood pressure is a major risk factor for coronary artery disease, heart failure, stroke, renal failure, and premature mortality. (Rapsomaniki et al., 2014) It is well-established that elevated low-density lipoprotein cholesterol (LDL-C) and low high-density lipoprotein cholesterol (HDL-C) are related to the future risk of cardiovascular disease (CVD) events. Wong et al. revealed that increased blood pressure (BP) confers increased risks for CVD in elderly persons across all lipid levels. These results document the importance of combined hypertension and dyslipidemia. (Wong et al., 2009). Kraja et al. identified SNP rs9678851 (missense) on gene SLC4A1AP associated with systolic blood pressure (SBP) in a meta analysis. SLC4A1AP is protein coding gene. It encodes a solute carrier also known as kidney anion exchanger adapter protein although it is widely expressed in most Genotype-Tissue Expression consortium tissues. (2017) Heart failure (HF) is a complex disease involving multiple changes including cardiomyocyte hypertrophy. Lu et al. (2012) performed a set of screens in different HF and hypertrophy models and identified gene SLC4A1AP as one the top ten differentially expressed genes associated with HF and/or hypertrophy. (Lu et al., 2012) In the genome-wide association Diabetes Genetics Initiative (D-GI) Study for 19 traits, including plasma lipids, SNP rs780094 in glucokinase (GCK) regulatory protein gene (GCKR) region was identified as a novel quantitative trait locus associated with plasma triglyceride concentration. (The Diabetes Genetics Initiative of the Broad Institute of MIT and Harvard, Lund University, and Novartis Institutes for BioMedical Research, 2007). GCKR rs780094 lies in a large region of linkage disequilibrium on chromosome 2. Both gene SUPT7L and SLC4A1AP are within the associated interval to be 417 kb based on linkage disequilibrium between the index SNP (rs780094) and SNPs upstream and downstream of the index SNP. (Orho-Melander et al., 2008). PTPN13 (also called FAP-1) is a non-receptor PTP and interacts with a number of important components of growth and apoptosis pathways. It is also involved in the inhibition of Fas-induced apoptosis. Laczmanska at el. (2017) identified an association of PTPN13 rs989902 with the risk of sporadic colorectal cancer in a Caucasian population. Adipose tissue plays critical roles in the regulation of energy homeostasis; as a reservoir, by storing and releasing fuel, and as an endocrine organ, by secreting a number of hormones and cytokines (Spiegelman and Flier, 2001). Excess body fat, or obesity, is a major public health problem increasing the risk of diabetes, cardiovascular diseases and several types of cancers (Aviva et al., 1999). Glondu-Lassis et al. (2008) demonstrated that knockdown of PTP-BL expression in 3T3-L1 adipocytes, a large cytoplasmic phosphatase also known as PTP-BAS/PTPN13/PTP-1E, caused a dramatic decrease in adipogenic gene expression levels and lipid accumulation. All these evidences suggest that these identified novel genes are likely to be associated with lipids or related traits. Further research is needed on the possible roles of these identified SNPs/genes.

Comparing the two real data analyses, our proposed GPA method can identify more significant genes. Specially, we can identify novel genes comparing with minimum p-values across all SNPs and our method is the most powerful test compared with other tests based on summary data. On the other hand, it has also been proven from the other side that SNPs do not work in isolation and a group of functionally related SNPs as annotated in a biological gene are often involved in the susceptibility and progression of the same disease. The results can be viewed as a complementary instead of competing approach to the traditional gene based test.

## Discussion

We have proposed gene-based adaptive combination method, which requires only p-value from GWAS summary data. With more and more GWAS summary data publicly accessible in the post-GWAS era, these methods based on summary data will be practically very useful to identify more genetic variants associated with various diseases. There are several important advantages of our method. First, it searches the most possible subset of SNPs by ranking p-values and identify strong association with gene avoiding the effect of other noises. Second, statistics for different types (e.g., dichotomous, ordinal, continuous) can all be easily analyzed by only using the p-values. Third, since our adaptive method is based on p-values from summary data for single trait, it can also be extended to test the association between multiple phenotypes and multiple genetic variants. Fourth, like other statistical methods based on summary data, the common feature is that they can increase sample size by combining summary statistics from a series of studies. Fifth, even though we usually can’t obtain distribution of the test statistics for many genetic association test methods, these methods can also be extended to perform summary data based test using learning from our permutation steps where these genetic association test methods can all be written as a function of Z-statistics. In conclusion, we showed that our proposed method provides a useful framework for gene-based association studies and a reference for other summary data based test methods.

As we know that expression quantitative trait loci (eQTLs) can affect complex traits by regulating gene expression levels and there is also an enrichment of eQTLs among the GWAS trait associated variants (Albert and Kruglyak, 2015; Lappalainen et al., 2013). Ignoring information on gene expression regulation may suffer loss of power for traditional gene-based tests. Thus, we note that we can readily integrate eQTL data (eQTL-derived weights) with summary data from GWAS to identify genes associated with a complex trait. Denote the gene expression derived weights as ***W*** = (*w*_1_,*…, w*_*M*_). Consider the weighted summary statistics based on Z-statistics (*w*_1_*Z*_1_,*…, w*_*M*_ *Z*_*M*_) and the corresponding associated covariance can be written as Σ(*σ*_*ij*_) : *σ*_*ij*_ = *w*_*i*_*r*_*ij*_*w*_*j*_. Specially, it will degenerate to traditional gene-based method when we set *w*_*m*_ = 1 for each *m*. The other settings for weights include 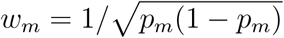,where *p*_*m*_ denote the MAF of the *m*^*th*^ variant (Madsen and Browning, 2009), 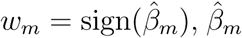 is an estimated value of the coefficient of the mth variant based on the marginal logistic model (Han and Pan, 2010) or *w*_*m*_ = *a*_*m*_*s*_*m*_*v*_*m*_ where *a*_*m*_ is a continuous weight (e.g., to incorporate allele frequencies), *s*_*m*_ determines the direction of the variant effect (deleterious or protective) and *v*_*m*_ is an indicator variable determining wether the variant belongs to the model (Hoffmann et al., 2010). We are currently exploring the weighted summary data based test methods and will report related results in the future.

The proposed method can be efficiently extended to analyze summary data for pathway-based analysis, because the pathway-based analysis has been proposed and applied in practice to boost statistical power and improve interpretability for GWAS (Pan, Kwak and Wei, 2015; Bakshi et al., 2016; Li et al., 2016). Therefore, we recommend applying the proposed test to further identify more novel genetic variants, especially for genes in the pathway with only small effect sizes. We have implemented the proposed methods in a C package available at https://github.com/Biocomputing-Research-Group/GPA. We provide sample C codes to install and use the package at the supplementary materials.

